# Rapid gastrointestinal tract transit time in volant vertebrates, with implications for gut microbiome convergence: Results from a meta-analysis

**DOI:** 10.1101/2024.08.09.607319

**Authors:** Emily Cornelius Ruhs, Katherine G. McFerrin, Devin N. Jones-Slobodian, Monique Ades, Radha Srinivasan, Natalia Cortes-Delgado, Ny Anjara Fifi Ravelomanantsoa, Carl J. Yeoman, Raina K. Plowright, Cara E. Brook

**Author notes:** Co-first author (Emily Cornelius Ruhs, Katherine G. McFerrin).

## Abstract

Flying birds and bats have simplified gastrointestinal tracts (GITs) and low intestinal mass to support flight. While previous work showed reduced GIT passage times in birds relative to other vertebrates, GIT passage times have never been collectively quantified for bats. We conducted a meta-analysis of published digesta passage times across vertebrates, comparing volant and non-volant vertebrates, while considering the effects of body mass and diet. We hypothesized that, like flying birds, bats have significantly faster digesta passage times relative to nonvolant vertebrates, likely due to their adaptations for flight. Our study supports this, revealing significant differences in passage times among flying and non-flying groups, with bats exhibiting faster transit times compared to non-volant vertebrates. Using a phylogenetic comparative analysis, we show that flight and diet have a strong effect on GIT transit times across diverse taxa, while body mass plays a more limited role. Accelerated transit times in bats likely promote the rapid turnover of gut contents, which may contribute to their distinct GIT microbe compositions. Unique among mammals, bat GIT microbiomes are dominated by Pseudomonadota bacteria, a pattern also observed in flying birds. We hypothesize this convergence may result from rapid GIT transit times quantified here for volant taxa.

## INTRODUCTION

Flying vertebrates (e.g., birds and bats) have unique and potentially convergent gastrointestinal physiologies, likely evolved to facilitate powered flight [1]. As one such adaptation, flying birds are known to have rapid gastrointestinal tract (GIT) passage times [2,3] (i.e., the time taken for food to pass through the stomach, small and large intestine, and colon prior to excretion [4]), a strategy hypothesized to help them maintain low digesta volume during flight [3]. The passage of digesta through the GIT is affected by many factors, several of which converge in flying birds and bats. In flying birds, rapid GIT transit is facilitated by short intestine lengths and elevated defecation and consumption rates to meet energy demands [2,5]. Bats share the short intestine lengths [6,7] and high defecation rates of flying birds [8,9]; in turn, faster digesta passage times also enable an increase in consumption rates during foraging [8]. Despite this, digesta passage times have never been collectively quantified for bats. We explore whether patterns of convergence in GIT physiology may be mirrored by convergence in digesta passage time in these two taxonomic groups.

Anatomically, longer and more complex GITs (e.g., multiple stomach chambers such as those found in herbivorous ungulates) tend to increase digesta passage time [10,11], whereas shorter, simpler GITs may decrease transit time [6,12]. Nonetheless, very few studies have quantified the correlation between GIT length (which scales allometrically with body size [13,14]) and transit time (but see [15–17]). As previously reviewed [12], relative to non-flying mammals, both small birds and bats have short, simple digestive tracts, with reduced nominal surface area and intestinal tissue [6,18]. Most bat GITs also lack a cecum and appendix, organs with specialized digestive properties in other taxa [18–21]. To compensate for the loss of intestinal absorptive surface area important for nutrient uptake in the GIT, both flying birds and bats have greater villous amplification (i.e., increase in digestive surface area) in the GIT epithelium and exhibit greater reliance on paracellular absorption (e.g., the passive uptake of water-soluble nutrients in the GIT) for meeting energetic demands [6,7].

In addition to anatomy, diet composition is correlated with differences in digesta passage times—though these differences may be most directly modulated by ingestion rate [22], which, in turn depends on nutrient composition (e.g., lower nutrient diets necessitate higher ingestion rates, which, consequently, yield more rapid GIT transit) [23]. Nonetheless, some evidence indicates that the water and fat content, consistency, and digestibility of ingested food can directly impact passage time [24,25]. For instance, insectivorous bats retain food longer than frugivorous bats, and some studies attribute this lengthier retention to a diet of more digestible food—though, notably, differences in intake volume may confound this relationship [26–28].

Evidence from pollen-eating bats indicates that the degree of dietary specialization is related to the efficiency of digestion as well: for example, flower-specialist bats digest pollen into usable protein resources more rapidly than generalist species [29,30]. Similarly, among insectivorous bats, the shortest GIT passage times have been recorded for trophic specialists [27]. It is also hypothesized that the microbiome may play a larger role in nutrient extraction for diet specialist bats and could explain some of the differences in the efficiency of digestion and gastrointestinal design, as well [31].

Interactions between the GIT microbial community and the host diet are modulated by digesta passage rates [12,32,33]. In humans, GIT transit is directly, positively correlated with total bacterial count [34,35] and positively associated with the presence of slow-growing gut microbes, such as methanogens [36,37]. In migratory bats, the core phyla of the GIT microbiome remain relatively stable despite major diet shifts due to long distance migration, possibly due to rapid digesta passage limiting microbiome restructuring [38]. Digesta passage times may also impact the capacity in which GIT microbes produce metabolites for their hosts. For example, short chain fatty acids (SCFAs) are produced through fiber breakdown by GIT microbes. SCFA production, in turn, stimulates intestinal motility for more rapid transit. High SCFA concentrations are also typically associated with lower GIT pH, which is associated with reduced transit time [33].

While most mammalian GIT microbiomes that have been surveyed to date (mainly those of Orders Artiodactyla, Carnivora, Primates, and Xenarthra) are primarily comprised of bacteria in the phyla Bacillota or Bacteroidota [39], both flying bird and bat GIT microbiomes are dominated by bacteria in the highly diverse phylum Pseudomonadota (previously Proteobacteria) [40–42]. This similarity of unusual gut microbiome composition between bats and birds suggests a possible shared mechanism; as Pseudomonadota are well known for their success in rapidly changing (and often aerobic) GIT environments [43–45], it is possible that similarly rapid GIT transit times in bats and flying birds may contribute to their proliferation in these two clades. Nonetheless, GIT transit has never been collectively summarized for bats.

In this study, we compiled GIT passage times from the literature for a wide range of volant (bats and flying-birds) and non-volant (carnivores, ungulates, rodents, primates, reptiles) vertebrate taxa. Using a phylogenetic comparative analysis, we tested the effects of flight, mass and diet on digesta passage times across taxa. We show that flying vertebrates (e.g., bats and birds) do indeed have significantly faster GIT passage times than non-volant vertebrates, offering one possible explanation for the congruent GIT microbiomes of bats and flying birds.

## METHODS

### Literature survey and statistical approach

We collected GIT passage times from the primary literature across a wide range of vertebrate taxa to assess the impact of flight on the duration of time required for food passage in the GIT. We constructed our dataset of GIT passage times by first conducting a strict keyword search for primary literature. First, we searched Web of Science for publications which cited the following search criteria in the title or abstract: “gut OR gastrointestinal OR food” AND “transit OR passage” AND “rate OR time”. To focus our study on non-model animal systems, we excluded articles that referenced human and laboratory mouse and/or rat studies using the “NOT” function for the following off-target terms: human[s], [wo]men, obesity, diabetes, adult, child[ren], patient[s], volunteer[s], disease, mice/mouse, rat[s]. This yielded 1169 studies.

We next individually reviewed each of the 1169 studies and retained any study (107) that reported “minimum gut transit” (MGT), “mean retention time” (MRT), “transit time”, or “gut passage time” for any vertebrate taxon in our final database. We then manually curated the database, adding 20 other published studies on digestive passage cross-referenced in the original database and incorporating 664 published works from the Abraham et al. 2021 dataset which were missed by the original search terms [46]. We examined each study to confirm that the value reported corresponded to a measure (e.g., transit time, MRT) of the duration of time between food consumption to first passage in feces, to ensure these values were comparable. Though MRT, the average period of time food is retained in the gut, offers an arguably more accurate representation of the digesta passage rates relevant to the establishment of GIT microbial communities [47], we found transit time (sometimes referenced as MGT) to be more widely reported in the literature across multiple vertebrate taxa and offer a larger pool of species for comparison than MRT. Thus, we report results for GIT transit time in the main analysis and results for MRT in the supplemental analysis, using the same statistical approach for both datasets (appendix 1). Of the studies reviewed, we identified 791 publications that reported GIT transit times for 243 species and MRT for 230 species for inclusion in our final database: (https://github.com/brooklabteam/git-transit-time).

Several factors may influence digesta passage time. Dry matter intake (DMI), or the quantity of food consumed, has been proposed as an influential predictor of digesta passage [48–50]. However, we did not include DMI in our analysis as these data were unavailable across many studies, and we aimed to prioritize species richness in our dataset. Despite our thorough review, we caveat that our database may have under- or overrepresented some taxa, particularly birds and small mammals, and it is possible that some species were missed.

From each study, we extracted the following information for each species surveyed and recorded it in the final database (Supplementary Dataset 1): (1) Order/Family/Genus/Species/Common Name, (2) body mass (grams), (3) food included in feeding trial (“experimental diet”, a broad description), (4) mean or median GIT transit time (or MRT) and, when available, minimum/maximum, and standard error/deviation of reported time, and finally (5) number of individuals and number of trials included in survey. We caveat that our conclusions may be limited by discrepancies between the experimental diet recorded in the trial and an organism’s natural diet outside captivity; however, manipulation of experimental diet was most likely to directly influence the GIT passage results surveyed here. We categorized experimental diets as unknown, fiber/foliage, fruit/nectar/pollen, meat, and protein. There were some animals (n=26 species) where the study [18] did not report the experimental diet; thus, diet was recorded as “unknown.” Experimental diets including dog or cat food, dairy, eggs, insects, and peanut butter were recorded as “protein,” whereas diets including beef, chicks, fish and mice were recorded as “meat”. If diets were reported as fish or meat/protein alongside fruits or fiber, they were recorded as “mixed”. In cases where a single study reported passage times for multiple individuals of the same species, we averaged the passage time per species reported from that study prior to further statistical analyses. When the transit time or MRT values were only reported in a figure, not as values in a table, we used PlotDigitizer (https://plotdigitizer.com) to extract data from graphs. When some of the species-level information (i.e., body mass) was missing from the original study, we searched the literature to find average values for the taxon in question. We recorded the source (if not the original study) and the method of data summarization for each entry in our database, adjacent to the metric in question. We removed two taxa (*Uromacer oxyrhynchus* and *Atheris squamigera)* because we could not find their masses reported in the literature.

We next compared GIT transit times across the different taxonomic groups surveyed. First, we used the unique transit time value for each entry in the database, preferentially selecting the mean reported value, then the median if mean was not reported. We then calculated an average transit time value for each species in our database, weighting database entries by the number of individuals surveyed and the number of trials conducted in each original study. When this original sample size information was not included in the study (n=52 entries), we assumed a sample size and trial size of only one individual in the weighting. We used the same approach to summarize average mass for each species reported in our database. Once the dataset was collapsed to a single transit time (and MRT when available) and mass for each species, we then compared digesta passage times across disparate taxa.

Due to small sample sizes, we elected to categorize all non-mammalian vertebrates taxonomically by Class (specifically, Birds, Reptiles, etc.) and all mammals by Order. We separated birds into “Flying” and “Non-Flying” categories, though transit time was only reported for five species of non-flying birds and MRT for seven species of non-flying birds. After assigning these groupings, we subset the database to include only those taxonomic groupings with more than four reported species and those with a reported experimental diet.

### Phylogenetic structure

To account for the possible role of phylogenetic relatedness in driving species-level patterns in transit time and MRT, we used the program TimeTree: The Timescale of Life [51] to construct a time-scaled phylogeny. All species represented in the cleaned database were available on TimeTree, except for n=5 species for transit time *(Adropadus milanjensis, Ceratogymna cylindricus, Cercopithecus albogularis schwarzi, Gulosus aristotelis, Eurillas latirostris)* and n=3 species for MRT (*Gazella subgutturosa marica, Gulosus aristotelis, Nectarinia famosa*). We removed these species from the final database for comparison (Table 1).

**Table 1.**
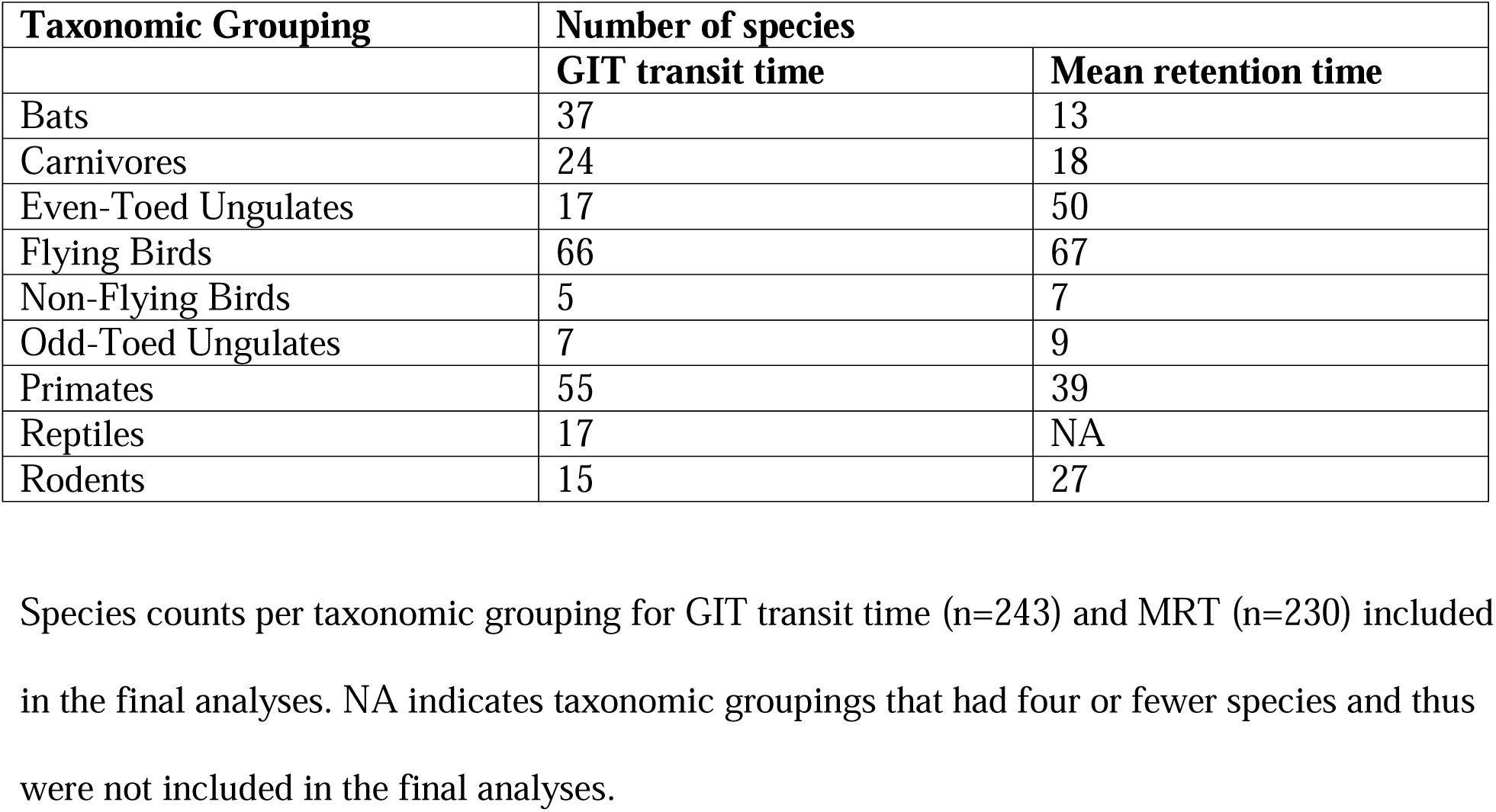
Species counts per taxonomic group for digesta passage metrics (GIT transit time and MRT).

| **Taxonomic Grouping** | **Number of species** |  |
|---|---|---|
|  | **GIT transit time** | **Mean retention time** |
| Bats | 37 | 13 |
| Carnivores | 24 | 18 |
| Even-Toed Ungulates | 17 | 50 |
| Flying Birds | 66 | 67 |
| Non-Flying Birds | 5 | 7 |
| Odd-Toed Ungulates | 7 | 9 |
| Primates | 55 | 39 |
| Reptiles | 17 | NA |
| Rodents | 15 | 27 |

### Phylogenetic signal

Before conducting our main analyses, we tested for a phylogenetic signal (phylosig) [52] for both transit time and MRT using the phytools package in R [53]. We quantified the strength of the phylogenetic signal by Pagel’s lambda, which can vary from no signal (0) to a strong signal (1) [46]. Pagel’s λ was first estimated from the full species tree and trait vector to quantify global phylogenetic signal. For both GIT and MRT, we compared models that fixed λ at 0 (no phylogenetic signal) against those that fixed λ at 1 (full phylogenetic signal) against those that fixed λ at the value estimated by phylosig for the corresponding dataset (λ=1.0015 for GIT; λ=0.82 for MRT). This approach ensured that all subsequent generalized least square (GLS) model comparisons were evaluated under an equivalent covariance structure; re estimating λ within each model would produce different phylogenetic correlation matrices between models, preventing direct Akaike Information Criterion (AIC) comparability. Because Pagel’s λ for GIT was equal (λ=1.0015) to the full phylogenetic model (λ=1), we compared only λ=0 against λ=1 models for these analyses. We conducted all subsequent model comparison using the nlme package in R [54,55].

### Model Structures and Comparisons: Predictors of Transit Time in Vertebrates

We constructed a series of candidate GLS models to test how flight, experimental diet, and body mass influence gastrointestinal transit time and mean retention time, while accounting for phylogenetic relatedness. All models were fit by maximum likelihood (ML) using nlme functions (gls, lme) [54]. For GIT transit, we compared model fits assuming λ=1 (full phylogenetic signal equivalent to Brownian motion) or λ=0 (no phylogenetic signal). For MRT, we compared model fits assuming λ=1, 0, or 0.82 (as estimated for our data by phylosig; see results).

We first tested for the effect of flight (flyer vs. non-flyer) on the response variables of transit time and MRT, without accounting for mass or diet, but comparing models accounting for phylogenetic structure (λ=1 and λ=0.82 for MRT) against those which did not (λ=0). We next asked whether inclusion of the additional predictor variable of experimental diet improved model fit to the data. For this second analysis, we compared structures modeling diet as a fixed or a random effect, again testing models accounting for phylogeny against those which did not. We next evaluated whether the addition of taxon mass as a fixed effect improved model fit to both GIT transit and MRT data, again comparing models accounting for phylogeny against those which did not. We considered disparate model structures accounting for mass and flight exclusively, in addition to those considering mass and flight simultaneously. Finally, we also tested whether mass alone (with no consideration of flight) was significantly predictive of transit time, both when accounting for phylogeny and otherwise.

In all cases, models were compared using AIC [56], with ΔAIC ≥ 2 considered support for selection of one model over another. In cases where models yielded equal support by AIC, we preferred the simplest model. For specific model structures see Table 2.

**Table 2.**
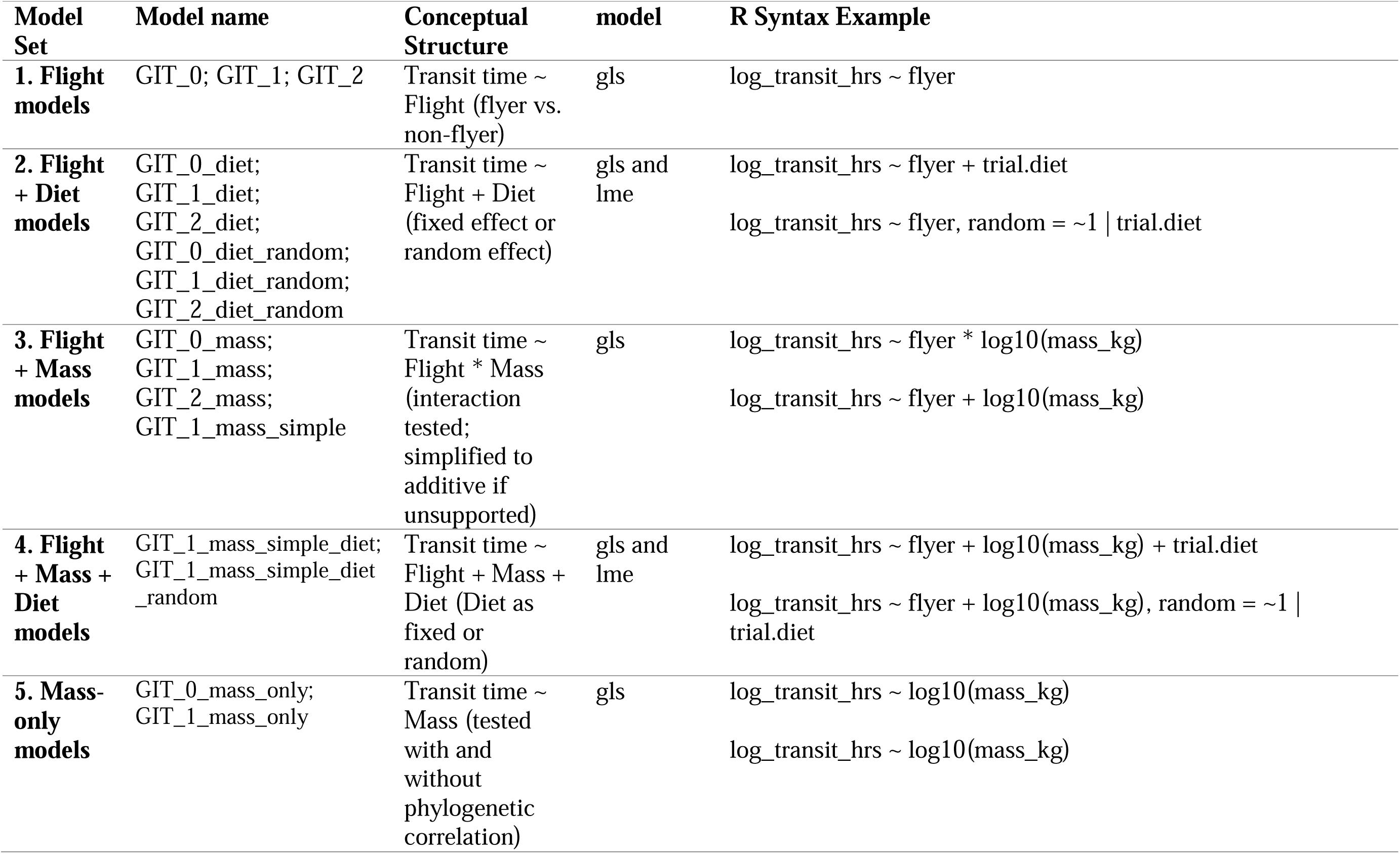
Outline of models included in our comparative analysis. Models were compared using both λ=1 and λ=0. For MRT, an additional lambda set by phylogenetic structure was used (λ=0.82).

## RESULTS

### Phylogenetic structure

We identified a significant phylogenetic signal in the GIT transit time data, at p<0.0001 with Pagel’s λ = 1.0015 (logL = −1553.37, LR = 151.226; Figure 1), and in the MRT data, at p<0.0001 with Pagel’s λ = 0.8185 (logL = −966.574, LR = 139.383; Figure S1). The higher estimate for Pagel’s λ indicated a stronger phylogenetic signal in the GIT transit vs. MRT data. Subsequently, we constructed models for GIT transit and MRT data which incorporated both λ= 1 and λ=0 and analyzed MRT data with an additional model that specified λ=0.82.

**Figure 1:**
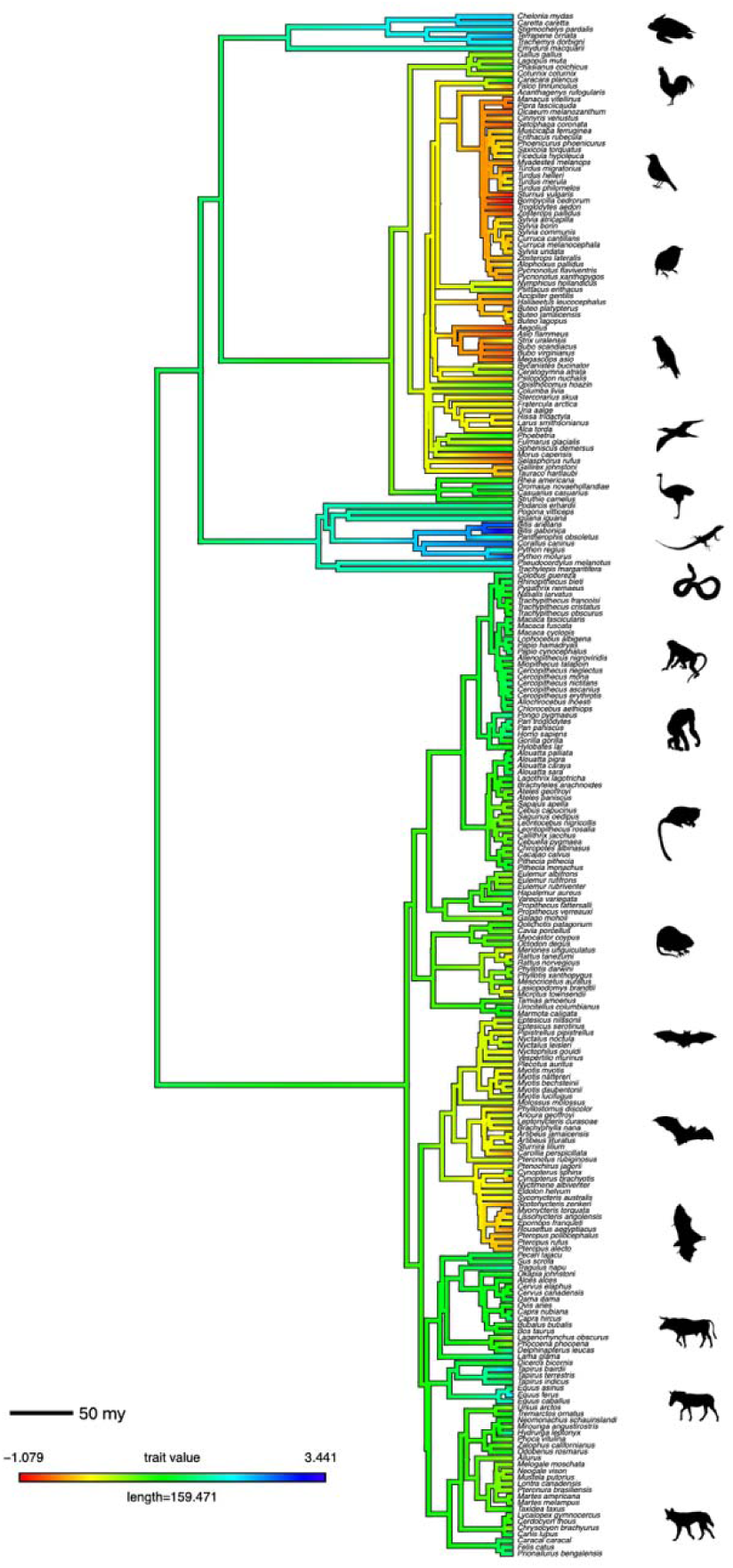
Phylogenetic mapping of GIT transit time across vertebrate taxa. A continuous trait map illustrating the evolution of log_10_-transformed GIT transit time across the phylogenetic tree of species included in our analysis. Branch colors represent reconstructed ancestral state trait values for relative GIT transit time, using a continuous gradient from slow (blue) to fast (red) transit times. The ancestral trait reconstruction was estimated using maximum likelihood, with tip labels corresponding to species names. The scale bar indicates relative branch length (in millions of years), and the color legend shows the range of trait values included in the reconstruction. Turtles (e.g., *Terrapene ornata*, *Caretta caretta*) and snakes (e.g., *Bitis gabonica*, *Python molurus)* show the slowest transit times, whereas bats (e.g., *Pteropus rufus*) and flying birds (e.g., *Troglodytes aedon*) show the fastest transit times.

### Comparative Predictors of GIT Transit Time in Vertebrates

Phylogenetic comparative analyses revealed strong effects of flight, mass, and diet on GIT transit time. Across all models, incorporating phylogenetic structure (λ = 1) consistently improved fit, as indicated by lower AIC values compared to models without phylogenetic structure (Table 3; Table S1).

**Table 3.** Model comparison results for GIT transit.

| Model name | Predictors | Phylogeny | df | AIC | Interpretation |
|---|---|---|---|---|---|
| GIT_0 | Flight | λ=0 | 3 | 468.54 | Flight is negatively associated with transit time (p<0.001), even accounting for phylogeny. |
| GIT_1 |  | λ=1 | 3 | -2281.11 |  |
| GIT_0_diet | Flight + Diet | λ=0 | 8 | 467.08 | Flight (p<0.001), diet unknown (p=0.02). |
| GIT_1_diet |  | λ=1 | 8 | -2287.23 |  |
| GIT_0_diet_random | Flight + 1\|Diet | λ=0 | 4 | 469.27 | Diet is better as a fixed effect |
| GIT_1_diet_random |  | λ=1 | 4 | -496.76 |  |
| GIT_0_mass | Flight*Mass | λ=0 | 5 | 465.64 | Flight significant (p<0.001), mass significant (p=0.001) |
| GIT_1_mass |  | λ=1 | 5 | -2288.44 |  |
| GIT_1_mass_simple | Flight + Mass | λ=1 | 4 | -2288.57 | Flight is significant (p<0.001). Mass also significant (p=0.002). |
| **GIT_1_mass_simple_diet** | **Flight + Mass + Diet** | **λ=1** | **9** | **-2295.67** | Diet is better as a fixed effect when mass is also considered |
| GIT_1_mass_simple_diet_random | Flight + Mass + 1\|Diet | λ=1 | 5 | -508.69 |  |
| GIT_0_mass_only | Mass | λ=0 | 3 | 610.00 | Mass is significant without phylogeny (p<0.001) |
| GIT_1_mass_only | Mass | λ=1 | 3 | -2261.71 | Mass is significant with phylogeny (p<0.001) |

In models that included flight as the sole fixed predictor, both with and without phylogenetic structure, flying species had significantly shorter transit times relative to non-flying taxa (p < 0.001 for both models; Figure 2a; Table S1). Because the effect of flight retained significance even after accounting for phylogeny, we concluded that flight itself—independent of phylogeny—has a significant negative relationship with GIT transit.

**Figure 2:**
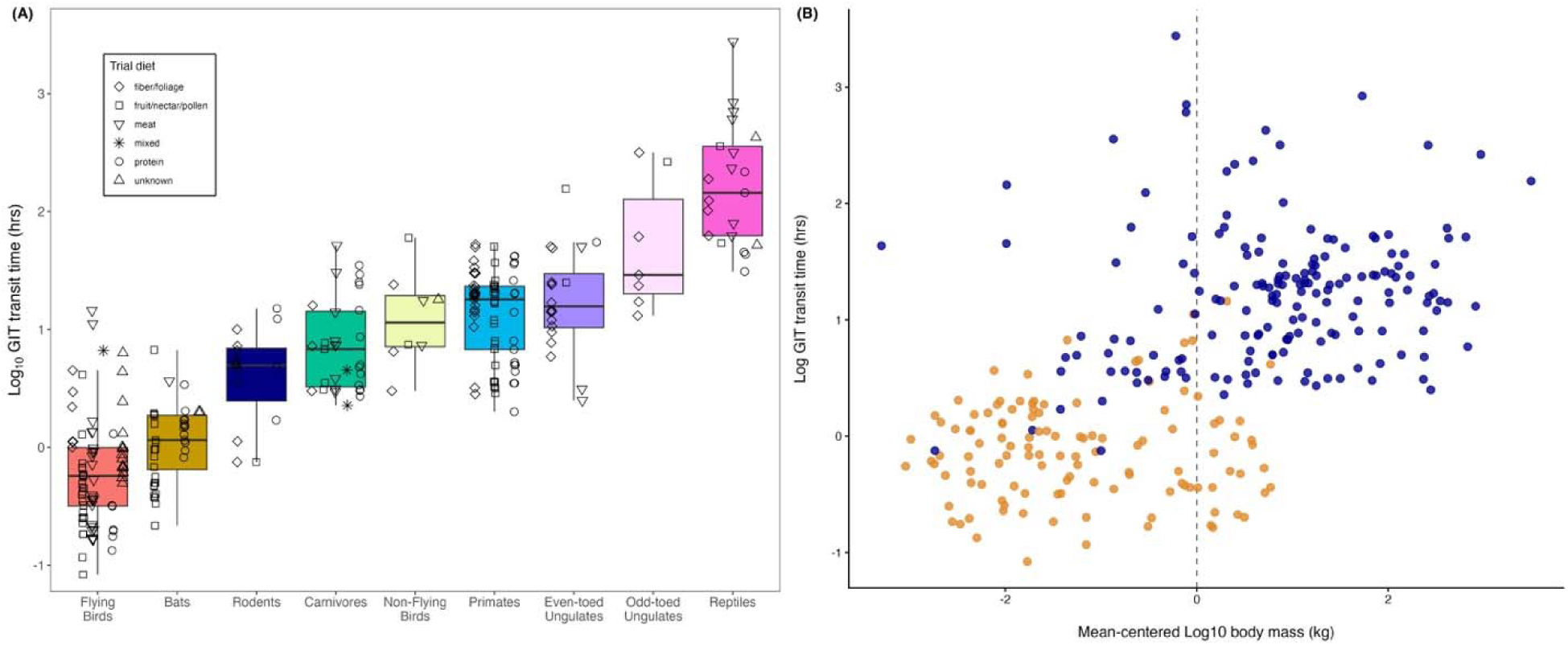
GIT transit times from experimental studies across volant and nonvolant vertebrates. **(a)** Boxplots show the interquartile range of GIT transit times (y-axis, log_10_ scale) recovered from studies across diverse vertebrate taxa (x-axis). Shapes correspond to experimental diet (according to the legend). Flying vertebrates (e.g., flying birds and bats) have significantly faster GIT transit times relative to all non-flying vertebrates (Table S1). **(b)** GIT transit time (y-axis, log_10_ scale) for flyers (orange) and non-flyers (blue) relative to their body mass (in kg; x-axis, log_10_ scale). Though Flight + Mass + Diet models offered the best fit to the GIT transit data, Flight + Diet and Flight + Mass models, also performed well (Table 3) and demonstrated a significant positive association between body mass and GIT transit time.

Incorporation of experimental diet as a fixed effect (but not as a random effect), in addition to flight, improved the fit of a phylogenetically-informed model over the flight-only construction (Table 3; Tables S1; Flight + Diet models). However, of the specified diets, only the ‘unknown’ category was significantly (positively) related to transit time (Table S1; p=0.02). The phylogenetically-informed model including body mass as a fixed effect (Flight + Mass model) additionally improved the fit of the Flight-only model (Table 3). However, the model which included fixed effects of flight, mass, and diet (without any interaction terms) provided the best fit overall (Table 3). Body mass demonstrated a significant positive correlation with GIT transit in Flight + Mass models (Table S1, p=0.002) and also was significant when also accounting for experimental diet (Flight + Mass + Diet models; p=0.001). In this latter case, the partial effects of each experimental diet retained largely the same significance as in Flight + Diet models not including mass. When we tested the effect of body mass on transit time without considering flight (Mass-only models), we found that body mass was a significant positive predictor of transit time regardless of the inclusion of phylogeny (Figure 2b; Table S1, p < 0.001 with and without phylogeny). Nonetheless, the significance of both flight and mass in all phylogenetically-informed models suggests an independent association of both features with GIT transit. In summary, our analyses identified flight as the most consistent significant predictor of reduced transit time, but models improved when diet as well as mass were additionally included fixed effects.

The phylogenetic distribution of GIT transit time further illustrates these trends (Figure 1). Flying clades, including birds and bats, consistently cluster toward lower trait values (e.g., faster transit times; orange-red branches), while non-flying mammals, birds, and reptiles exhibit longer transit times (e.g., slower transit times; green-blue branches).

### Comparison of digesta passage time metrics: GIT transit time and MRT

Across both metrics of digesta passage, GIT transit time and MRT, the best supported models were phylogenetically informed and included mass or flight as a key predictor. However, in contrast to GIT transit models, the phylogenetically-informed mass-only model offered a competitive fit to the MRT data, as compared with Flight + Mass and Flight + Mass + diet models (Table S2). In all models tested, body mass had a significant positive effect on both GIT transit time and MRT (Table S1, Table S2, Table S3, p <0.001), irrespective of whether phylogeny was accounted for. In contrast to GIT transit time analyses, flight was only a marginally significant (negative) correlate of MRT in the phylogenetically-informed Flight + Mass or Flight + Mass + diet fixed effect models (Table S3). These results suggested that some of negative effects of flight on digesta passage time (specifically, MRT) may be explained by the small average body masses of bats and birds. Interestingly, in the Flight + Mass models with interaction terms, the interaction of flight with mass had a marginally significant positive association with MRT (Table S3, p=0.09), suggesting that larger fliers may have longer MRTs. Taken together, both GIT transit time and MRT analyses demonstrate a significant association between smaller, flying vertebrate taxa (bats and birds) and faster digesta passage than that exhibited by larger, non-flying vertebrates.

## DISCUSSION

Understanding the factors influencing digesta passage times across vertebrate taxa is crucial for unraveling the intricate relationship between digestive physiology, diet, and evolutionary adaptations [46,48]. In this study, we aimed to compare digesta passage times among volant (bats and flying birds) and non-volant vertebrates while considering the effects of flight, experimental diet, body mass, and phylogenetic structure. Our findings shed light on the unique digestive adaptations of flying mammals – i.e., bats – and draw attention to the impacts that these adaptations may have on GIT microbial communities.

Consistent with previous research [10,16], our study reveals significant differences in digesta passage times among flying vertebrate groups. In our analyses, flight was the most influential predictor of digesta passage time for the GIT transit dataset, with flying species demonstrating rapid digesta passage compared to non-flyers, after controlling for effects of phylogeny and experimental diet. Including experimental diet as a fixed effect improved model performance over flight-only models for GIT transit time, though no particular experimental diet demonstrated any clear relationship with GIT transit. For the MRT data, inclusion of diet did not significantly improve model performance in any case. Because flight imposes exceptionally high metabolic demands, birds and bats have evolved to process food swiftly and extract energy-rich compounds with great efficiency. Rapid passage times in flying vertebrate taxa may also reflect high defecation frequencies through which quickly clearing undigested waste (i.e. feces) minimizes excess body mass to better support highly mobile lifestyles [9,57].

We found that, in addition to flight, body mass also demonstrated a significant effect on digesta passage time, in both GIT transit and, particularly, MRT datasets. Generally, larger-bodied species exhibited longer transit times, consistent with some allometric scaling theory equating larger mass and longer gastrointestinal tracts [13]. However, the effects of body mass on GIT transit and MRT showed a consistently positive relationship regardless of phylogenetic signal or the inclusion of flight in model structure, offering support for some universally predictable allometric scaling of GIT length with body size [46,48,58]. Nonetheless, because flight + mass models outperformed mass-only models in GIT transit analyses and offered comparable fits to the data for MRT analyses, our results suggest that volant species, regardless of size, consistently exhibit reduced baseline transit times relative to non-volant species. In general, flying vertebrates demonstrate both reduced body mass and shorter intestines compared to most non-volant mammals [3,14], which we here show to be associated with rapid GIT transit. This pattern indicates an ecological trade-off imposed by flight, as rapid GIT transit likely reduces opportunities for nutrient uptake in flighted taxa and may contribute to patterns of convergence in microbiome composition across flying vertebrates. This hypothesis is supported by the dominance of Pseudomonadota in volant bats and birds, in contrast to non-flying taxa (including flightless birds) [42]. Future research should explore the functional significance of GIT microbiome composition in flying vertebrates, with implications for both host nutrient acquisition and immunity.

## Supporting information

Supplemental analysis

## DECLARATIONS

### IRB statement

Not applicable.

### Author’s contributions

ECR conducted the final data analysis and wrote the first draft of manuscript. KGM collected data from the literature, helped with the data analysis, and shared in the writing of the first draft of manuscript. DNJS conceptualized the project, collected the original data from the literature, and contributed to writing of the manuscript. MA and RS helped with data curation. NCD reviewed the data and contributed to writing of the manuscript. NAFR helped with the data analysis and collection. CJY helped with the data analysis and writing. RKP conceptualized the project and provided funding. CEB conceptualized the project, collected the original data, helped with data analysis, funded the project, and contributed to writing of the manuscript. All authors reviewed and provided edits to the final manuscript.

## Acknowledgements

The authors thank the Brook lab at the University of Chicago for helpful comments on the manuscript.

## Availability of data and material

The datasets generated and the code used for analysis for the present study are available at: https://github.com/brooklabteam/git-transit-time.git

## Funding

The authors acknowledge funding support from a Branco Weiss ‘Society in Science’ fellowship to CEB, an NIH-NIAD DP2 award to CEB (1DP2AI171120), a pilot grant from the Microbiome Center at the University of Chicago to CEB, the DARPA PREEMPT Program Cooperative Agreement #D18AC00031 to RPK (with a subgrant to CEB), and a U.S. National Science Foundation grant (DEB-1716698, EF-2133763/EF-2231624) to RKP. CJY was funded through the Montana Agricultural Experiment Station (MONB00113). ECR was funded by a Grainger Bioinformatics Fellowship through the Field Museum of Natural History (Chicago, IL), and NCD was funded by a Chicago Fellows scholarship through the Biological Sciences Division of the University of Chicago. The content of the information does not necessarily reflect the position or the policy of the U.S. government, and no official endorsement should be inferred.

## Notes

https://github.com/brooklabteam/git-transit-time.git

### Competing Interest Statement

The authors have declared no competing interest.

### Summary of Updates

We have added additional peer-reviewed publications to our meta-analysis, consistent with reviewer comments and requests.

