## Supplemental analysis for "Rapid gastrointestinal tract transit time in volant vertebrates, with implications for gut microbiome convergence: Results from a meta-analysis"

**Supplementary Text: *Comparative Predictors of Mean Retention Time in Vertebrates***

Phylogenetic comparative analyses revealed strong effects of flight on mean retention time (MRT). Across all models, incorporating full phylogenetic signal equivalent to Brownian motion (λ = 1) consistently improved fit, compared to λ = 0.82 (as estimated by phylosig evaluation of the data), or compared to models without phylogenetic control (Table S2).

The best-supported models included flight as a key predictor, with flying species showing significantly shorter MRTs relative to non-flying taxa, after accounting for phylogenetic signal (Table S3, p < 0.001). In contrast to GIT transit models (main text), models that additionally incorporated diet as either a fixed or random effect did not improve model fit to MRT data after accounting for flight and phylogeny (Table S2). In Flight + Diet models for MRT, of the specified experimental diets, only the meat diet was significantly (positively) associated with MRT (Table S3; p=0.004).

When not accounting for flight, body mass alone was strongly positively correlated with MRT when phylogeny was not considered (Table S3, mass-only models, λ = 0; p<0.001) as well as in both phylogenetically-informed models (Table S3, mass-only models, λ = 1 or λ = 0.82). In combined models (Table S3, flight + mass and flight + mass + diet models), mass was always a significant positive predictor of MRT; additionally, the addition of mass to the base flight-only or flight + diet Model significantly improved model fit to the MRT data (Table S2).

Again, similar to GIT transit time, the phylogenetic distribution of retention time further illustrates these trends (Figure S1). Flying clades (e.g., bats), consistently cluster toward faster transit times (e.g. orange-red branches), while non-flying birds, reptiles, and mammals exhibit longer transit times (e.g. green-blue branches).

**Table S1. Model comparison outputs with effect sizes, 95% confidence intervals, and p-values (GIT transit time data).**

| **Model name** | **Phylogeny** | **AIC** | **Predictors** | **Effect size** | **SE** | **CI (LL)** | **CI (UL)** | **p-value** |
| --- | --- | --- | --- | --- | --- | --- | --- | --- |
| GIT_0 | λ=0 | 468.54 | Flight | -1.32 | 0.06 | -1.44 | -1.20 | <0.001 |
| GIT_1 | λ=1 | -2281.11 | Flight | -1.12 | 0.15 | -1.41 | -0.83 | <0.001 |
| GIT_0_diet | λ=0 | 467.08 | Flight | -1.36 | 0.07 | -1.50 | -1.22 | <0.001 |
|  |  |  | Fruit/nectar/pollen | -0.11 | 0.09 | -0.29 | 0.07 | 0.21 |
|  |  |  | Meat | 0.09 | 0.1 | -0.11 | 0.29 | 0.37 |
|  |  |  | Mixed | -0.17 | 0.32 | -0.80 | 0.46 | 0.59 |
|  |  |  | Protein | -0.09 | 0.1 | -0.29 | 0.11 | 0.37 |
|  |  |  | Unknown | 0.25 | 0.14 | -0.02 | 0.52 | 0.08 |
| GIT_1_diet | λ=1 | -2287.23 | Flight | -1.12 | 0.15 | -1.41 | -0.83 | <0.001 |
|  |  |  | Fruit/nectar/pollen | -0.11 | 0.07 | -0.25 | 0.03 | 0.14 |
|  |  |  | Meat | -0.02 | 0.10 | -0.22 | 0.18 | 0.82 |
|  |  |  | Mixed | -0.17 | 0.23 | -0.62 | 0.28 | 0.46 |
|  |  |  | Protein | -0.11 | 0.08 | -0.27 | 0.05 | 0.18 |
|  |  |  | Unknown | 0.27 | 0.12 | 0.03 | 0.51 | 0.02 |
| GIT_0_diet_random | λ=0 | 469.27 | Flight | -1.33 | 0.07 | -1.47 | -1.19 | <0.001 |
| GIT_1_diet_random | λ=1 | -496.76 | Flight | -1.05 | 0.16 | -1.36 | -0.74 | <0.001 |
| GIT_0_mass | λ=0 | 465.64 | Log_10_(mass) | 0.09 | 0.04 | 0.01 | 0.17 | 0.01 |
|  |  |  | Flight | -1.26 | 0.11 | -1.48 | -1.04 | <0.001 |
|  |  |  | Log_10_(mass):flight | -0.07 | 0.06 | -0.19 | 0.05 | 0.02 |
| GIT_1_mass | λ=1 | -2288.44 | Log_10_(mass) | 0.13 | 0.04 | 0.05 | 0.21 | 0.001 |
|  |  |  | Flight | -0.99 | 0.18 | -1.34 | -0.64 | <0.001 |
|  |  |  | Log_10_(mass):flight | -0.09 | 0.07 | -0.23 | 0.05 | 0.17 |
| GIT_1_mass_simple | λ=1 | -2288.57 | Flight | -0.91 | 0.17 | -1.38 | -0.44 | <0.001 |
|  |  |  | Log_10_(mass) | 0.10 | 0.03 | 0.04 | 0.16 | 0.002 |
| **GIT_1_mass_simple_diet** | **λ=1** | **-2295.67** | **Flight** | **-0.91** | **0.17** | -1.24 | -0.58 | **<0.001** |
|  |  |  | **Log_10_(mass)** | **0.10** | **0.03** | 0.04 | 0.16 | **0.001** |
|  |  |  | **Fruit/nectar/pollen** | **-0.08** | **0.07** | -0.22 | 0.06 | **0.29** |
|  |  |  | **Meat** | **-0.08** | **0.10** | -0.28 | 0.12 | **0.43** |
|  |  |  | **Mixed** | **-0.20** | **0.23** | -0.65 | 0.25 | **0.39** |
|  |  |  | **Protein** | **-0.07** | **0.08** | -0.23 | 0.09 | **0.35** |
|  |  |  | **Unknown** | **0.31** | **0.11** | 0.09 | 0.53 | **0.008** |
| GIT_1_mass_simple_diet  _random | λ=1 | -508.69 | Flight | -0.81 | 0.17 | -1.14 | -0.48 | <0.001 |
|  |  |  | Log_10_(mass) | 0.11 | 0.03 | 0.05 | 0.17 | <0.001 |
| GIT_0_mass_only | λ=0 | 610.00 | Log_10_(mass) | 0.33 | 0.03 | 0.28 | 0.40 | <0.001 |
| GIT_1_mass_only | λ=1 | -2261.71 | Log_10_(mass) | 0.17 | 0.03 | 0.11 | 0.23 | <0.001 |

**Table S2. Model comparison results for mean retention time (MRT).** In addition to setting lambda to 0 and 1, we included an additional lambda set by the data at (0.82; models with “set”).

| **Model name** | **Predictors** | **Phylogeny** | **df** | **AIC** | **Interpretation** |
| --- | --- | --- | --- | --- | --- |
| MRT_0 | Flight | λ=0 | 3 | 505.76 | Flight is negatively associated with MRT (p<0.001), even accounting for phylogeny. Assuming full Brownian motion is best supported. |
| MRT_set |  | λ=0.82 | 3 | -2951.90 |  |
| MRT_1 |  | λ=1 | 3 | -3331.37 |  |
| MRT_0_diet | Flight + Diet | λ=0 | 8 | 495.88 | Flight is still negatively associated with MRT (p=0.0004); however, only meat diet type is significant. |
| MRT_set_diet |  | λ=0.82 |  | -2954.12 |  |
| MRT_1_diet |  | λ=1 |  | -3330.50 |  |
| MRT_0_diet_random | Flight + 1\|Diet | λ=0 | 4 | 501.86 | Diet is better as a fixed effect |
| MRT_set_diet_random |  | λ=0.82 | 4 | -1253.91 |  |
| MRT_1_diet_random |  | λ=1 | 4 | -1544.88 |  |
| MRT_0_mass | **Flight*Mass** | λ=0 | 5 | 479.52 | **Flight is not significantly associated with MRT when mass and interaction terms are included (p=0.4), thought flight:mass interaction is marginally significant (p=0.09).** |
| MRT_set_mass |  | λ=0.82 | 5 | -2961.98 |  |
| **MRT_1_mass** |  | **λ=1** | **5** | **-3350.2** |  |
| **MRT_1_mass_simple** | **Flight + Mass** | **λ=1** | **4** | **-3349.35** | **Mass is significant (p<0.001). Flight is only marginally significant (p=0.12).** |
| **MRT_1_mass_simple_diet** | **Flight + Mass + Diet** | **λ=1** | **10** | -3347.66 | **Mass is significant (p<0.001). Flight is marginally significant (p=0.09).** |
| MRT_1_mass_simple_diet  _random | Flight + Mass + 1\|Diet | λ=1 | 5 | -1551.14 | Not supported |
| MRT_0_mass_only | Mass | λ=0 | 3 | 535.01 | **Mass was significant for all models and positively associated with MRT.** |
| MRT_set_mass_only | Mass | λ=0.82 | 3 | -2961.76  **-3348.96** |  |
| **MRT_1_mass_only** | **Mass** | **λ=1** | **3** |  |  |

**Table S3. Model comparison outputs with effect sizes, 95% confidence intervals, and p-values (MRT data).**

| **Model name** | **Phylogeny** | **AIC** | **Predictors** | **Effect size** | **SE** | **CI (LL)** | **CI (UL)** | **p-value** |
| --- | --- | --- | --- | --- | --- | --- | --- | --- |
| MRT_0 | λ=0 | 505.76 | Flight | -0.99 | 0.07 | -1.13 | -0.85 | <0.001 |
| MRT_1 | λ=1 | -3331.37 | Flight | -0.68 | 0.19 | -1.05 | -0.31 | <0.001 |
| MRT_2 | λ=0.82 | -2951.9 | Flight | -0.63 | 0.16 | -0.94 | -0.32 | <0.001 |
| MRT_0_diet | λ=0 | 495.88 | Flight | -1.0 | 0.07 | -1.14 | -0.86 | <0.001 |
|  |  |  | Fruit/nectar/pollen | -0.18 | 0.09 | -0.36 | 0.00 | 0.03 |
|  |  |  | Meat | 0.28 | 0.11 | 0.06 | 0.50 | 0.009 |
|  |  |  | Mixed | -0.27 | 0.39 | -1.03 | 0.49 | 0.49 |
|  |  |  | Protein | -0.06 | 0.11 | -0.28 | 0.16 | 0.57 |
|  |  |  | Unknown | 0.47 | 0.25 | -0.02 | 0.96 | 0.06 |
| MRT_1_diet | λ=1 | -3330.5 | Flight | -0.68 | 0.19 | -1.05 | -0.31 | <0.001 |
|  |  |  | Fruit/nectar/pollen | 0.03 | 0.09 | -0.15 | 0.21 | 0.74 |
|  |  |  | Meat | 0.39 | 0.14 | 0.12 | 0.66 | 0.004 |
|  |  |  | Mixed | 0.09 | 0.38 | -0.65 | 0.83 | 0.81 |
|  |  |  | Protein | 0.039 | 0.12 | -0.20 | 0.27 | 0.74 |
|  |  |  | Unknown | 0.07 | 0.2 | -0.32 | 0.46 | 0.71 |
| MRT_2_diet | λ=0.82 | -2954.12 | Flight | -0.64 | 0.16 | -0.95 | -0.33 | 0.001 |
|  |  |  | Fruit/nectar/pollen | 0.04 | 0.08 | -0.12 | 0.20 | 0.62 |
|  |  |  | Meat | 0.4 | 0.12 | 0.16 | 0.64 | 0.001 |
|  |  |  | Mixed | 0.1 | 0.35 | -0.59 | 0.79 | 0.78 |
|  |  |  | Protein | 0.04 | 0.11 | -0.20 | 0.28 | 0.71 |
|  |  |  | Unknown | 0.23 | 0.2 | -0.16 | 0.62 | 0.25 |
| MRT_0_diet_random | λ=0 | 501.86 | Flight | -1.0 | 0.07 | -1.14 | -0.86 | <0.001 |
| MRT_1_diet_random | λ=1 | -1544.88 | Flight | -0.65 | 0.16 | -0.96 | -0.34 | <0.001 |
| MRT_2_diet_random | λ=0.82 | -1253.91 | Flight | -0.66 | 0.14 | -0.92 | -0.38 | <0.001 |
| MRT_0_mass | λ=0 | 479.52 | Log_10_(mass) | 0.09 | 0.03 | 0.03 | 0.15 | 0.006 |
|  |  |  | Flight | -0.49 | 0.11 | -0.71 | -0.27 | <0.001 |
|  |  |  | Log_10_(mass):flight | 0.19 | 0.07 | 0.05 | 0.33 | 0.004 |
| **MRT_1_mass** | **λ=1** | **-3350.2** | **Log_10_(mass)** | **0.14** | **0.05** | 0.04 | 0.24 | **0.004** |
|  |  |  | **Flight** | **-0.17** | **0.22** | -0.60 | 0.26 | **0.44** |
|  |  |  | **Log_10_(mass):flight** | **0.15** | **0.09** | -0.03 | 0.33 | **0.09** |
| MRT_2_mass | λ=0.82 | -2961.98 | Log_10_(mass) | 0.12 | 0.04 | 0.04 | 0.20 | 0.007 |
|  |  |  | Flight | -0.28 | 0.2 | -0.67 | 0.11 | 0.17 |
|  |  |  | Log_10_(mass):flight | 0.07 | 0.09 | -0.11 | 0.25 | 0.44 |
| **MRT_1_mass_simple** | **λ=1** | **-3349.35** | **Flight** | **-0.31** | **0.2** | -0.70 | 0.08 | **0.12** |
|  |  |  | **Log_10_(mass)** | **0.18** | **0.04** | 0.10 | 0.26 | **<0.001** |
| MRT_1_mass_simple_diet | λ=1 | -3347.66 | Flight | -0.34 | 0.2 | -0.73 | 0.05 | 0.09 |
|  |  |  | Log_10_(mass) | 0.18 | 0.04 | 0.10 | 0.26 | <0.001 |
|  |  |  | Fruit/nectar/pollen | 0.05 | 0.08 | -0.11 | 0.21 | 0.52 |
|  |  |  | Meat | 0.35 | 0.13 | 0.10 | 0.60 | 0.008 |
|  |  |  | Mixed | 0.15 | 0.37 | -0.58 | 0.88 | 0.68 |
|  |  |  | Protein | 0.15 | 0.12 | -0.09 | 0.39 | 0.2 |
|  |  |  | Unknown | 0.14 | 0.19 | -0.23 | 0.51 | 0.46 |
| MRT_1_mass_simple_diet  _random | λ=1 | -1551.14 | Flight | -0.47 | 0.17 | -0.80 | -0.14 | 0.006 |
|  |  |  | Log_10_(mass) | 0.09 | 0.03 | 0.03 | 0.15 | 0.004 |
| MRT_0_mass_only | λ=0 | 535.01 | Log_10_(mass) | 0.28 | 0.02 | 0.24 | 0.32 | <0.001 |
| **MRT_1_mass_only** | **λ=1** | **-3348.96** | **Log_10_(mass)** | **0.2** | **0.04** | 0.12 | 0.28 | **<0.001** |
| MRT_2_mass_only | λ=0.82 | -2961.76 | Log_10_(mass) | 0.17 | 0.03 | 0.11 | 0.23 | <0.001 |

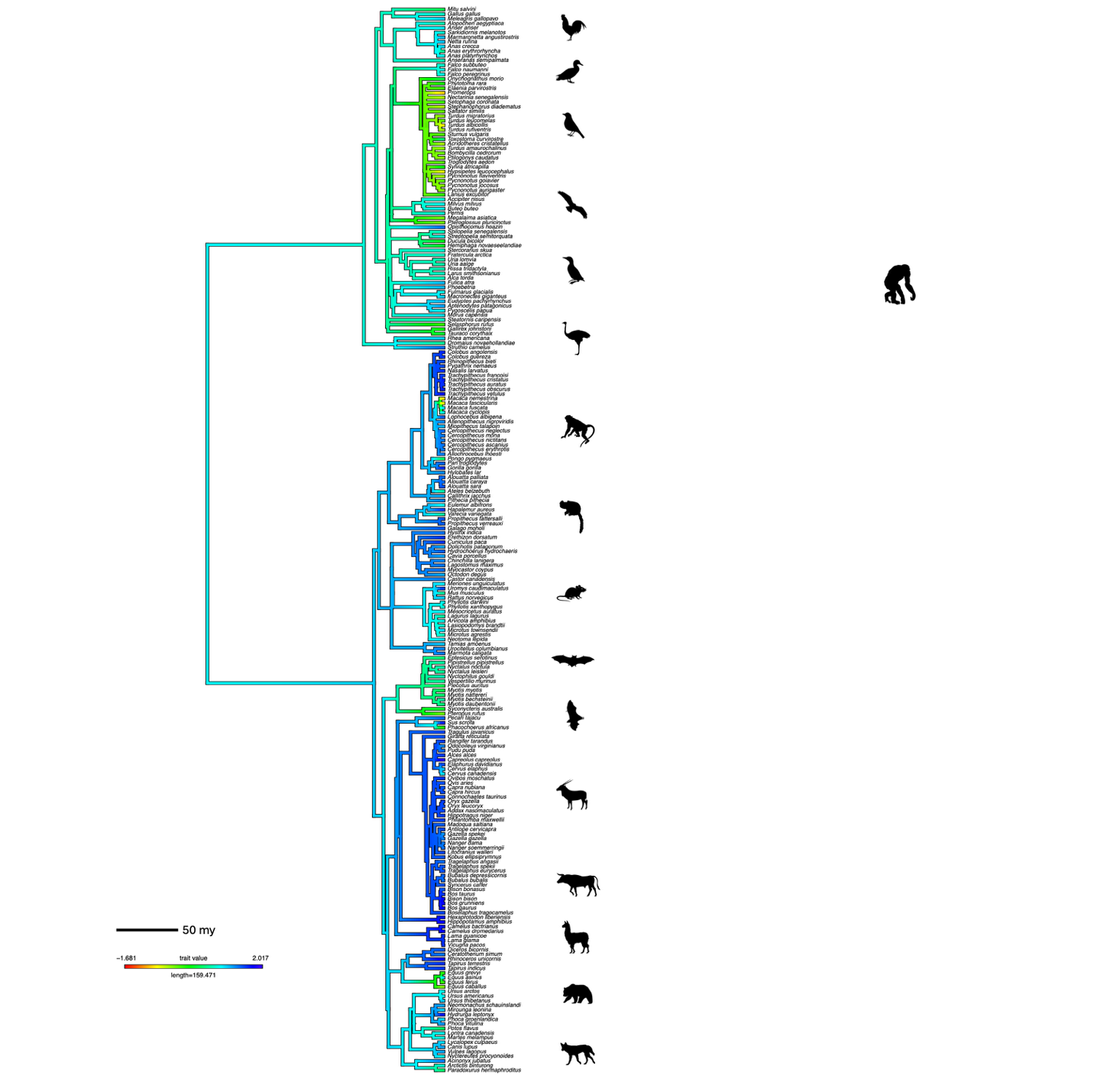

**Figure S1:** *Phylogenetic mapping of mean retention time (MRT) across vertebrate taxa.* A continuous trait map illustrating the evolution of log_10_-transformed MRT across the phylogenetic tree of species included in our analysis. Branch colors represent reconstructed ancestral state trait values for relative MRT using a continuous gradient from slower (blue) to faster (yellow) MRT. The ancestral trait reconstruction was estimated using maximum likelihood, with tip labels corresponding to species names. The scale bar indicates relative branch length (in millions of years), and the color legend shows the range of trait values included in the reconstruction. The ungulates (e.g. *Capra hircus, Bos taurus*) and some primates (e.g. *Pongo pygmaeus*) show the slowest MRTs, whereas bats (e.g., *Pteropus rufus*) and flying-birds (e.g., *Turdus albicollis*) show the fastest MRTs.

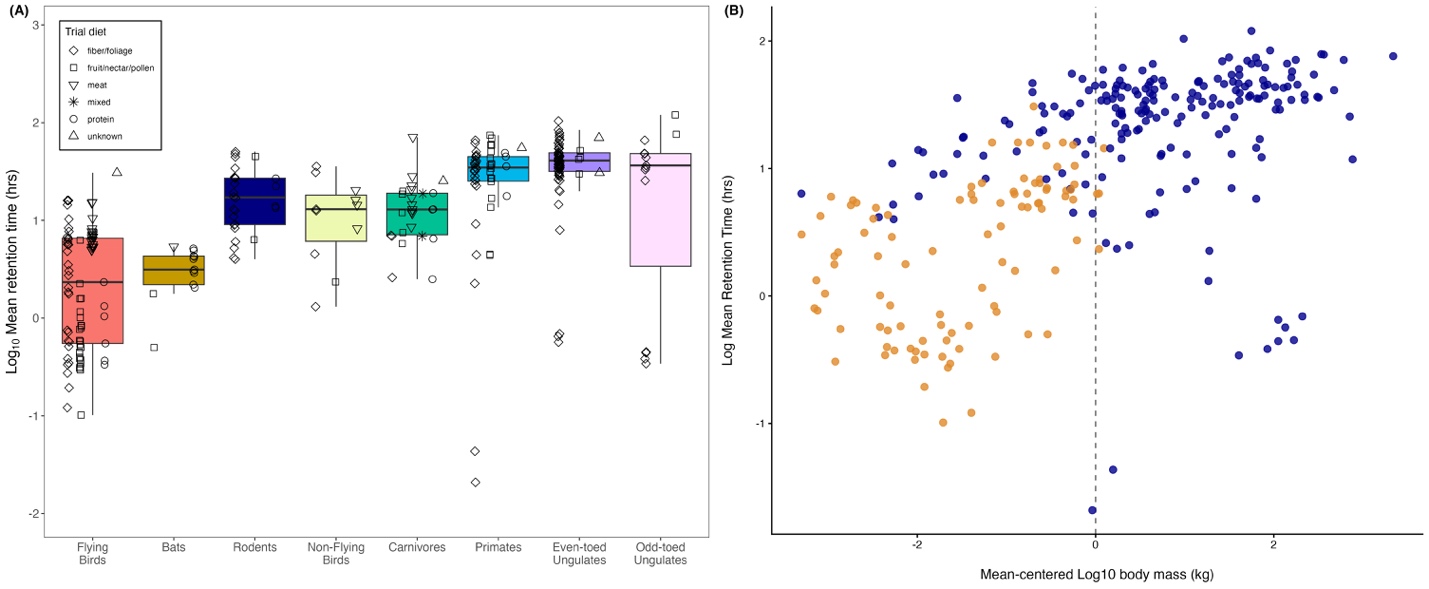

**Figure S2**: *Mean retention times (MRTs) from experimental studies across volant and nonvolant vertebrates*. **(a)** Boxplots show the interquartile range of MRTs (y-axis, log_10_ scale) recovered from studies across diverse vertebrate taxa (x-axis). Shapes correspond to food type typically consumed by each taxon (experimental diet, according to the legend). Flying vertebrates (e.g., birds and bats) have significantly faster MRTs relative to non-flying vertebrates in the analysis. **(b)** MRT (y-axis, log_10_ scale) for flyers (orange) and non-flyers (blue) relative to their body mass (in kg; x-axis). Body mass was a significant (positive) predictor of MRT in Flight + Mass, Flight + Mass + Diet and Mass-only models, which offered similarly competitive fits to the MRT data (Table S2, Table S3).
